# Rubisco Dark Inhibition in Angiosperms Shows a Complex Distribution Pattern

**DOI:** 10.1101/2025.11.20.689527

**Authors:** Connor Nehls-Ramos, Elizabete Carmo-Silva, Douglas J. Orr

## Abstract

Crop yields can be improved through making photosynthesis more efficient. The regulation of the CO_2_-fixing enzyme Rubisco during fluctuating light conditions limits productivity and is a target for improvement. Regulation in low light and darkness by accumulation of Rubisco inhibitors, predominantly via 2-carboxy-d-arabinitol 1-phosphate, has been known for over 4 decades but an explanation is still lacking for its physiological role and high variability across species. We compiled all published data for dark inhibition of Rubisco in flowering plants and investigated phylogenetic trends. Literature data for 157 species across 14 orders was compared and standardised, categorised into four dark inhibition levels, and analysed in the context of current phylogenetic information. We created a novel resource for Rubisco dark inhibition across flowering plants, highlighting clear gaps and biases in the available data, while also raising further questions on the evolution of this trait. Our work supports better understanding of the enigmatic process of photosynthetic regulation by Rubisco dark and low light inhibition and informs future efforts in enhancing photosynthesis in crops.

**Highlight:** Compilation and analysis of all available data on Rubisco dark inhibition from literature to date showed a complex distribution of levels of inhibition of the CO_2_ fixing enzyme in flowering plants. Such a distribution suggests there may be chloroplast microenvironment drivers for low *vs.* high Rubisco inhibition.

## Introduction

As global food insecurity increases, there is an ever-greater need for higher crop yield to contend with both losses by extreme climate events and meet growing demands (FSIN and GNAC 2025). One strategy to improve crop output is by improving photosynthesis through fixing inefficiencies associated with Rubisco, the imperfect enzyme essential to carbon fixation in photosynthesis (Croce et al. 2024). Despite advances in understanding the complex regulation of Rubisco activity, there remain many large knowledge gaps (Parry et al. 2008; Leister et al. 2023; Orr et al. 2023; Pasch et al. 2024). Passing clouds, wind, and sunflecks result in irregularities in lighting throughout the crop canopy. For many but not all crops, accumulation of Rubisco inhibitors has been reported when leaves experience an extended low light or dark period.

The identification of dark and low light Rubisco inhibition arose first due to anomalies in measuring the activation state of Rubisco in leaf extracts (Vu et al. 1984). Total activity measurements – incubating leaf extracts in an activating buffer containing Mg^2+^ and CO_2_ – showed a stark depression in activity in predawn samples compared to midday leaves in certain species. The *in vitro* carbamylation of Rubisco enhanced binding of dark and low light accumulated inhibitory compounds present in the leaf extracts, a process generally referred to as dark inhibition (Servaites 1985; Holbrook et al. 1992). The accumulation of inhibitors in the chloroplast is a slow process requiring hours in full darkness to reach peak inhibitor levels (Andralojc et al. 1994). In species with especially high dark inhibition of Rubisco, for example French bean (*Phaseolus vulgaris*) and potato (*Solanum tuberosum*), 2-carboxy-d-arabinitol 1-phosphate (CA1P) could explain the majority activity loss in the dark (Seemann et al. 1985; Gutteridge et al. 1986; Berry et al. 1987). Interestingly, as more species were studied for dark inhibition, it became clearer some plants lacked any appreciable accumulation of dark inhibitors (Holbrook et al. 1992).

Dark inhibition varies greatly between flowering plant species and, to a lesser extent, within varieties of the same species (Servaites et al. 1986; Holbrook et al. 1992; Sage 1993; Holbrook et al.1994; Orr et al. 2023). While the pathway for the *in vivo* synthesis of CA1P has yet to be conclusively resolved, the precursor molecule 2-carboxyarabinitol is suggested to be phosphorylated by a kinase with activity stimulated in low light (Andralojc et al. 1996a; Andralojc et al. 1996b; Andralojc et al. 2002; Parry et al. 2008). Regularly utilizing a kinase is resource and energy costly, implying an advantage to the accumulation of CA1P in the dark and low light, i.e., this inhibitor may provide benefits for plants depending on various environmental and metabolic pressures. Alternatively, CA1P accumulation may be a byproduct of a yet unknown metabolic pathway not harmful enough to be selected out by certain environments. Various hypotheses exist for the utility of dark inhibition or CA1P accumulation, but no conclusive *in vivo* evidence has yet been provided (Andralojc et al. 1996a; Khan et al. 1999; Van Heerden et al. 2003; Parry et al. 2012). The high variability in levels of accumulation, even across closely related species, suggests further nuance in the evolution and potentially the function of this trait.

While previous works have discussed trends in dark inhibition by phylogeny, no comprehensive list has been made across flowering plants (Holbrook et al. 1992; Sage 1993; Orr et al. 2023). To better understand the natural variation in dark inhibition, we have compiled the averaged species dark inhibition data from available literature for 157 species across 14 orders and organised this data by phylogeny and levels of dark inhibition. Data was also analysed and filtered by experimental methods to ensure data used were comparable and then summarized at species levels. Trends in dark inhibition level by phylogeny, photosynthetic metabolism type, and gaps in dark inhibition data from species to order level, suggest distinct order to species level evolutionary patterns in flowering plants and that dark inhibition may be older than flowering plants.

## Materials and methods

### Dark Inhibition Database

Dark inhibition data was collected from an extensive search of literature published, as of August 2025, using a combination of Google Scholar (Google 2025), Web of Science (Clarivate 2025), Scopus (Elsevier 2025), and a further search in the references of all papers found that contained information pertaining to Rubisco activity in light- and dark-adapted leaf samples. The keywords used for these searches were: “Rubisco”, “Dark”, “Inhibition”, “Diurnal”, “Total Activity”, “CA1P”, and “2-carboxy-D-arabinitol 1-phosphate”. These keywords were used in various combinations to maximize paper identification. The focus of these analyses is within flowering plants and so three species outside this criterion were not included for further analysis (see dataset). In total, 45 sources were found with 331 data points for dark inhibition. Each datapoint is representative of a given genotype sampled within each given study, or of the same genotype at different stages of growth. The full dataset is within the GitHub repository https://github.com/cwnehls/NehlsRamos_etal_JxB_BriefComms_Code_and_Dataset in the excel file NehlsRamos_etal_JxB_BriefComms_Data.

The dark inhibition ratio was calculated as [1 - (dark-adapted Rubisco total activity / light-adapted Rubisco total activity)]. For dark inhibition levels, this ratio is multiplied by 100 to represent the percentage of dark inhibition. Total Rubisco activity is the *in vitro* carboxylation activity for fully carbamylated Rubisco from leaf samples. For sources that did not include total activity values but had total activity plotted, the data was estimated using SplineCloud plot digitizer (Splinecloud 2025). Further details on light levels, literature sources, and more are contained in the dark inhibition dataset excel file.

Species phylogenetic information was collected using the angiosperm phylogeny website, NCBI taxonomy browser, The Kew Tree of Life, and the most recent literature on Plant taxonomic groupings (Ruggiero et al. 2015; Baker et al. 2022; Stevens 2025; Sayers et al. 2025).

Data quality assessment for final analysis included preliminary comparison of variation at species level of the three main experimental methods used for activity determination in the literature collected, 3PGA-NADH coupled assays, radiolabelled CA1P assays, radiolabelled CO_2_ assays and (Supplementary Fig. S1a). Results were inconsistent for the first two assay methods for species dark inhibition values, compared with the radiolabelled CO_2_ method. While the 3PGA-NADH method did not have significant differences for the proportion of dark inhibition between dark and light treated samples in, such as in wheat (Sales et al. 2020), the lack of data with these methods provides insufficient information to more conclusively determine if the results from these methods are comparable to the radiolabelled method that has been used for the large majority of species. To minimize unnecessary unknown variation while retaining the highest amount of quality data, only the radiolabelled CO_2_ method was kept for further analysis.

### Radiolabelled-Method Data Validation

Analysis was performed for the difference in reagent concentration, assay timing, plant sampling conditions, as well as pH and temperature of assays. This did not identify significant trends by linear regression, meta regression and Kruskal-Wallis tests, besides the light level for dark treatments and EDTA concentrations (data not shown). Slight differences in methods, such as addition of MgCl_2_ in the sample extraction buffer (Sharwood et al. 2016) or purity of RuBP (Andralojc et al. 2012), can have significant effects on recorded total Rubisco activity. However, in the context of the proportion of activity from light- and dark-treated samples, these discrepancies did not provide significant statistically identifiable differences that could otherwise be explained by species level variation (Supplementary Fig. S1b; see code). Furthermore, at species level for *Phaseolus vulgaris*, the most sampled species, a Kruskal-Wallis test detected a significant but relatively small difference with light level of sampling. In *Glycine max*, the next most sampled species, this difference was not found. It was concluded that there was insufficient evidence for variation within the radiolabelled CO_2_ method applications that would justify removal of specific studies, and so all datapoints for flowering plants using these methods were kept for further analysis. The final count for literature sources was 39, with 312 data points on dark inhibition, 157 of which are from unique species (see full dataset).

### Dark Inhibition Levels

By comparison of normal distribution, gaussian fit, and data point clustering, four main categories of dark inhibition were identified within the current data. The distribution of the data at order level was determined to be not normal by Hartigan’s Dip Test. To visualize if there were clustering of dark inhibition values a combination of Gaussian Mixture Models by McLust, Silverman’s tests for modes (data not shown), and Kernel Density Estimate modelling showed four distinct peaks within the distribution of species dark inhibition values (Supplementary Fig. S2a). The antimodes, or valleys, between these modal peaks remained similar with and without the presence of the Fabales data, which was tested to ensure overrepresentation of Fabales did not skew the categorisations (Supplementary Fig. 2b, c). These levels of dark inhibition are described as low (<18%), moderate (18-44%), High (44-77%), and very high (>77%) (Supplementary Fig. S2A). While high variation of dark inhibition values was found at even species level for some species (Table 1), the presence of the four modes provided a range of values that could be used to describe the general distribution of dark inhibition at species, genus, or order level for identification of further trends.

**Table 1.**
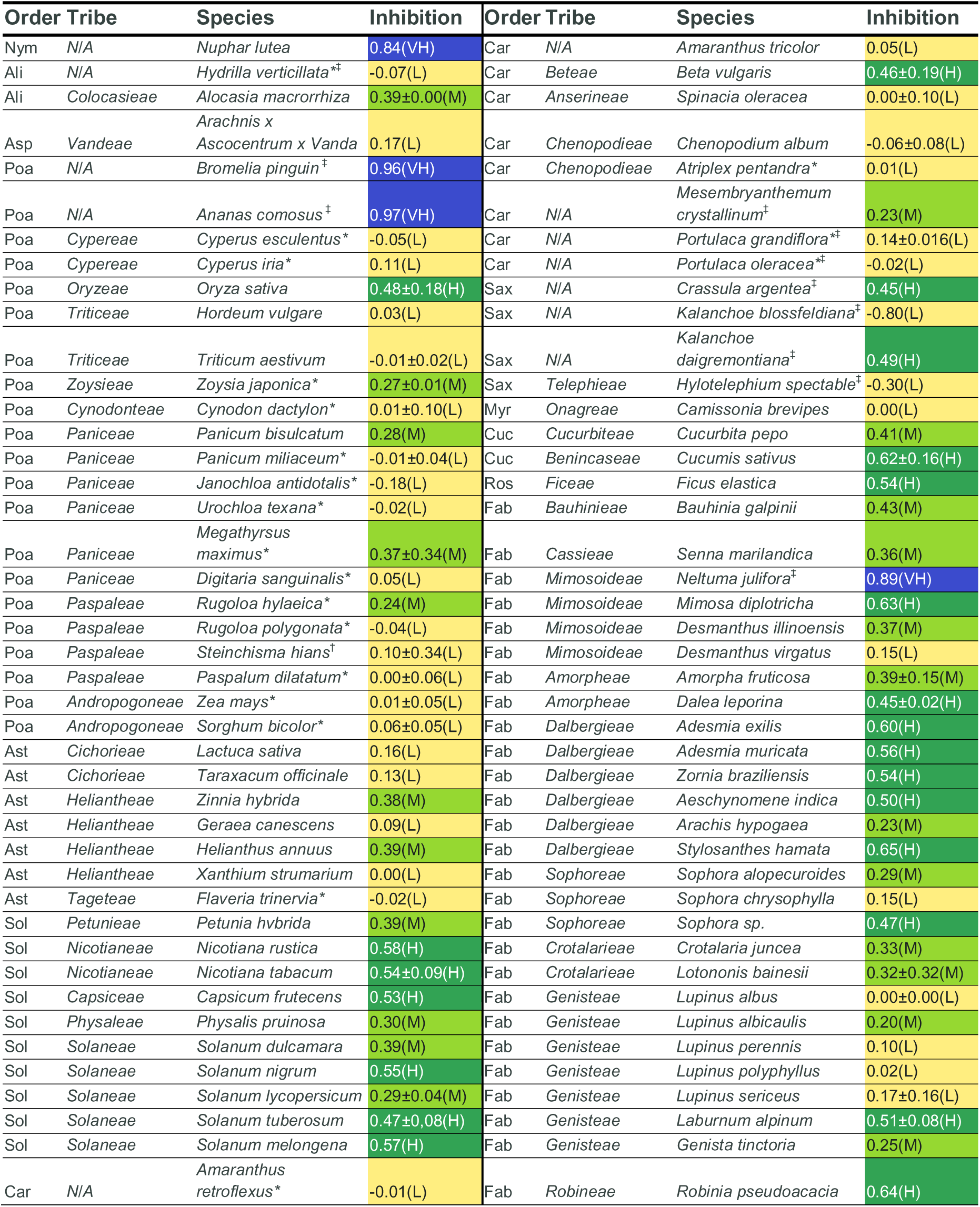
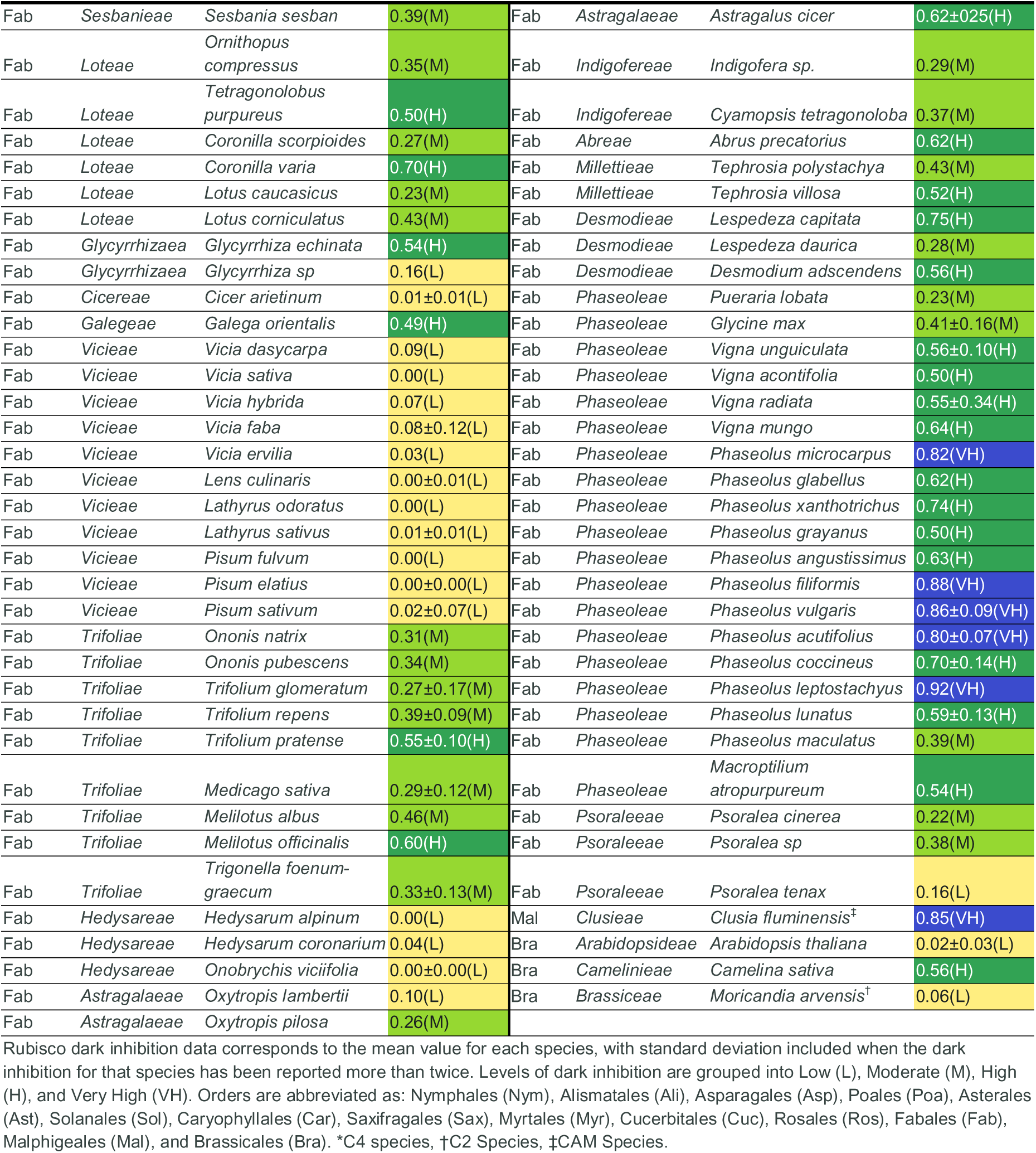
Rubisco dark inhibition measurements for each species to date by order and tribe.

### Data Visualisation

Microsoft Excel was used to visualise data in Figure 1a and Table 1. And R was used to produce Figure 2, using the following packages: ggplot2; multcompView; dplyr; readxl; agricolae; tibble; forcats; car; FSA; rcompanion; viridisLite; metafor; clubSandwich; robumeta; pracma; diptest; mclust; multimode. Figure 1b was created through using the Kew Tree of life species tree as the backbone, labelling and colouring branches and nodes using the Interactive Tree of Life website, then further labelling the image in PowerPoint (Baker et al. 2022, Letunic and Bork 2024). The coloured lines signifying average genera dark inhibition are for all monophyletic species in the tree of the given genera, even if only one species was studied for that genus. The tree can be accessed publicly here: (https://itol.embl.de/tree/148882475326681755591567). All analyses and data are available in GitHub (<u>cwnehls/NehlsRamos_etal_JxB_BriefComms_Code_and_Dataset: Code for use in the paper, Nehls-Ramos et al. 2025. See readme for more information</u>).

**Fig. 1.**
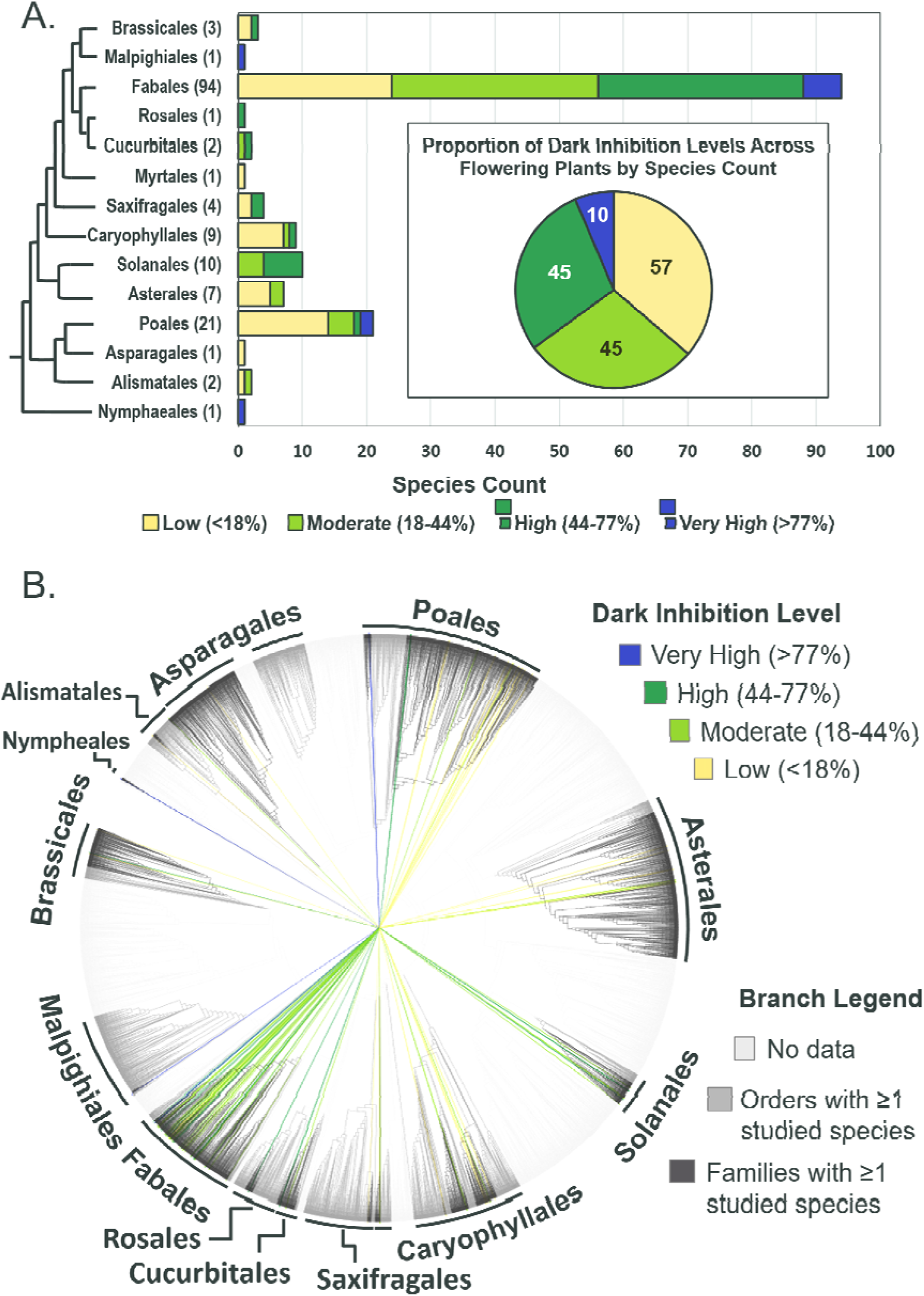
Distribution of Rubisco dark inhibition levels across flowering plant orders. (A) Simplified tree showing the phylogenetic relationship of fourteen orders represented in the literature. Stacked bars represent the average species dark inhibition level, grouped from low, moderate, high and very high. In parentheses is the number of unique species studied within each order. Inset: breakdown of proportion of species dark inhibition across all species studied, numbered by count. (B) Phylogenetic tree of over 10,000 flowering plant species coloured by dark inhibition levels at genera level. Branches are coloured light grey for no species data available, grey for orders with one or more species data available, and dark grey for families with one or more species data on dark inhibition available. Lines radiating from the centre of the tree are connect to respective genera and coloured to indicate the average dark inhibition level of that genera. Dark inhibition levels are coloured yellow for low (<18%), yellow-green for moderate (18-44%), green for high (44-77%), and blue for very high (>77%).

**Fig. 2.**
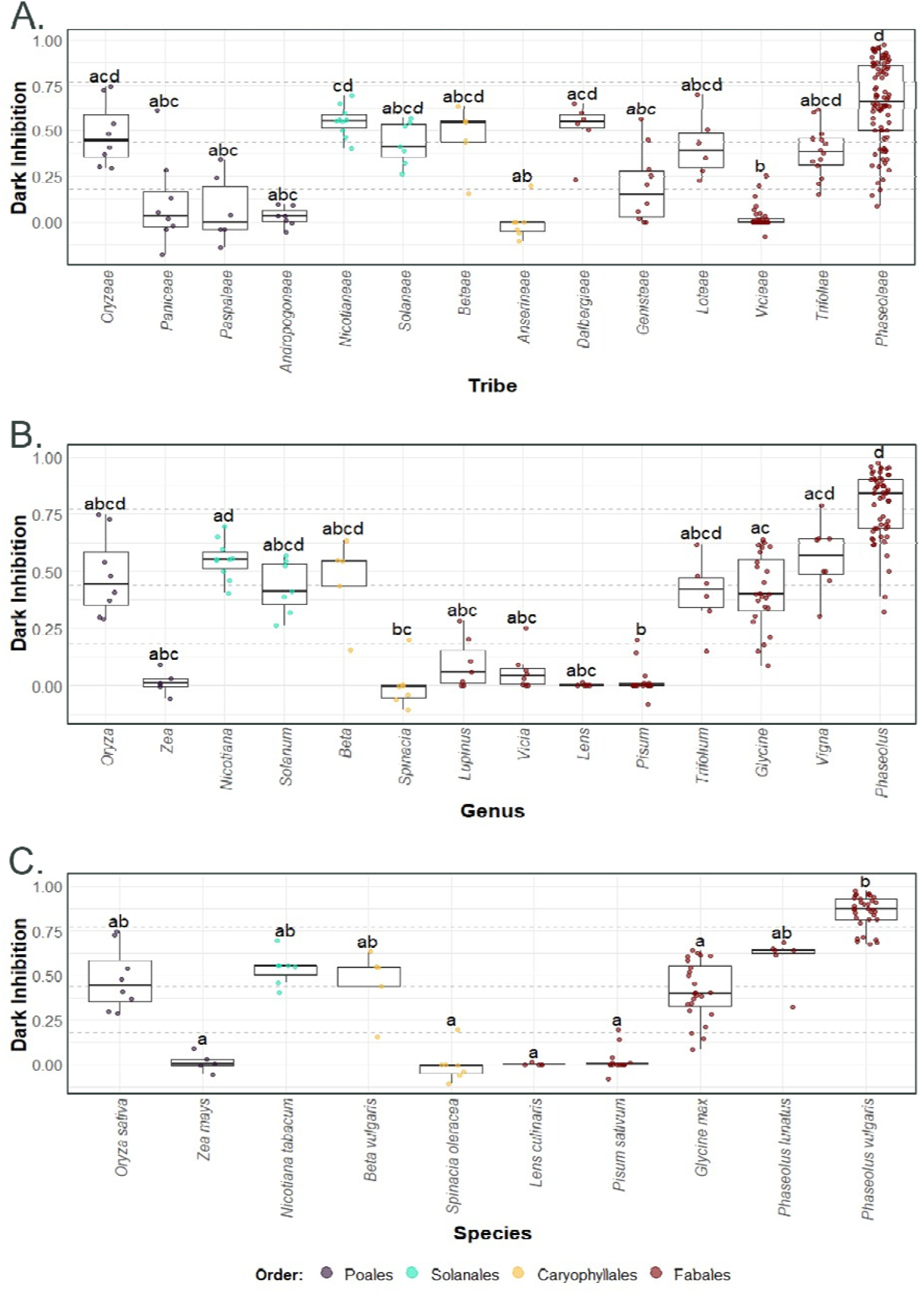
Variation in Rubisco dark inhibition distribution at different taxonomic levels. (A) Tribe, (B) Genera, and (C) Species. Only groups with a minimum of five data points are included. Data points are colored by order, purple is Poales, teal is Solanes, yellow is Caryophyllales, and dark red is Fabales. All groups are ordered by phylogenetic proximity. Dark inhibition ranges from 0 to 1, indicating 0% to 100% inhibition respectively. Box plots show medians and the first and third quartiles (25^th^ and 75^th^ percentiles), and whiskers extend from the hinge to the largest or smallest value. Kruskal-Wallis followed by Dunn *post hoc* tests were performed to identify significant differences between groups, denoted by different letters (*P*<0.05).

### Data Analyses

Kruskal-Wallis and meta regression modelling was used to assess whether data on Rubisco dark inhibition were broadly comparable between different Rubisco activity methods and methodological variations of these, as well as different publications. One-way ANOVA was not possible due to the consistent lack of similar sample size between groups, and lack of normality as determined by Shapiro-wilk test. The full dataset for flowering plants obtained with studies employing the radiolabelled CO_2_ method was found to not be normally distributed and had unequal variance. Determination of Rubisco dark inhibition levels to establish groups of low to very high inhibition used a combination of McLust Gaussian mixture models, Hartigan dip test, followed by kernel density estimates to determine number of modes in the data, and subsequently the position of modes and antimodes. For comparative analyses of dark inhibition across phylogenetic groups, Kruskal-Wallis followed by Dunn *post hoc* tests, with the Holm adjustment, were used due to the dataset not being normally distributed, the sample sizes often being too small, and showing uneven variance. All tests assumed a 0.05 significance level.

## Results

The dark inhibition data across studies carried a high sampling bias for the order Fabales, with far less sampling data regarding most other studied orders. Fabales, with 94 of the 157 species, represented approximately 60% of the species level data. Poales, with 21 species, was the only other order for which more than 10 species have had Rubisco dark inhibition data reported to date (Fig. 1a). This sampling bias is explained by two studies which contribute numerous species, all within the order Fabales (Holbrook 1992; Sage 1993). From the data available we identified different levels of dark inhibition across species and provide a comprehensive survey on dark inhibition at species level, which presented interesting trends in distribution across flowering plants (Fig. 1 and Table 1).

Proportional differences in species dark inhibition levels were visible across orders. Dark inhibition was classified into four levels, low (<18%), moderate (18-44%), high (44-77%), and very high (>77%) (See Materials and Methods). Focusing on the two most studied orders, Poales showed a greater proportion of low dark inhibition levels compared to the more even distribution across inhibition levels in Fabales (Fig. 1a). Grouping species dark inhibition levels by phylogenetic order showed no clear trends across orders, though some had distinct trends even at low species count-- for example Solanales, with 10 species studied, had only moderate and high inhibition levels. Nonetheless, across all studied species low inhibition was the most common level (57 of 157 species), being highly prevalent within Poales and Caryophyllales. Moderate and high levels were the next highest (45 species each). Very high inhibition levels were the rarest, with <7% of all species studied (10), half of these being members of Fabales.

To visualise further phylogenetic trends, distribution of these levels was compared across a comprehensive phylogenetic tree for flowering plants. The categorised dark inhibition levels were overlaid across the Kew Tree of Life with branch tips at species level and labelling at genus level (Fig. 1b). The data available to date represents 14 of 64 orders of flowering plants, with wide coverage across the phylogeny tree. Within the gaps are smaller crop relevant orders and families containing numerous spices, fruits, vegetables, and nuts. From current data, the more basal Poales branches containing the bromeliads and *Oryza* have higher dark inhibition compared to derived Poaceae grass lineages (Fig. 1b). Within Fabales, the highly studied Fabaceae family holds a wide spread of moderate and high inhibition genera, with low inhibition genera sequestered in distinct branches, while the *Phaseolus* branch trends towards very high inhibition (Fig. 1b).

Patterns at tribe level and below were explored using a comprehensive survey on dark inhibition at species level (Table 1). While nearly a quarter of Fabales species have low dark inhibition, these species are concentrated in the genera *Lupinus, Cicer, Hedysarum*, and *Onobrychis,* and the tribe *Viceae*. In contrast, other branches of Fabales, such as *Phaseolus* and *Vigna,* show above average Rubisco dark inhibition, with *Phaseolus* containing the most ‘very high’ level species of all studied genera in flowering plants (Table 1). Within this genus is *P. vulgaris*, the most studied species for dark inhibition. *Glycine max*, a close relative of *Phaseolus* and *Vigna,* had lower comparative mean inhibition and the highest intraspecies variation (±14%) for well-studied species-- the driver for this surprising variation is discussed later (Table 1).

Dark inhibition values showed inconsistent levels of variation at both genus and species level across flowering plants. Comparing groups with 5 or observations there were significant differences at tribe level, with seven tribes showing lower Rubisco dark inhibition levels compared with *Phaseolae* (Fig. 2a). At order level, significant differences were observed between the moderate and high inhibition Fabales and Solanales and the low inhibition Poales and Caryophyllales (Supplementary Fig. 3a). At genus level, this trend was also observed in relation to *Phaseolus* (Fig. 2b). This was observed despite the high range of variation in *Phaseolus* with a median of 80% dark inhibition and a standard deviation of more than 20%. There were eight genera with moderate to high inhibition and in some cases significant differences were observed between these and the low inhibition genera *Pisum* and *Spinacia* (Fig. 2b). At species level, significant differences were only identified for low versus the very high inhibition species, *P. vulgaris* (Fig. 2c). Some taxa showed trends for lower or higher inhibition that were not significant, potentially due to low sample size resulting in insufficient statistical power. Conversely, for the plants with more data availability, there were clear and statistically significant differences. This was particularly the case for Fabales, with some genera showing near-zero dark inhibition compared to high levels observed in others, especially *Phaseolus* (Supplementary Fig. S3b).

## Discussion

Rubisco dark inhibition data published to date was compiled and analysed to identify trends that would help better understand the underlying role of this regulatory mechanism. Across flowering plants, only 157 species have been measured for Rubisco dark inhibition, representing less than 0.05% of the more than 300,000 known species, and only 14 of 64 orders (Baker et al. 2022; Antonelli et al. 2023). The data available shows bias towards the Fabales order, consisting of nearly two thirds of the total species measured with wide dark inhibition variation across species.

Intraspecies variation observed may potentially be due to biological variation, differences in sampling, plant age, growth conditions, assay methods, as well as a variable presence of daytime inhibitors (Vu et al. 1983; Moore et al. 1995). While factors such as assay methods and conditions were investigated here, for others there is insufficient data currently available. Cultivar differences were visible in *G. max,* even within the same study, though current data make robust comparisons of cultivar level variation difficult (Holbrook et al. 1994). Species level variance was far less present in related beans, *Vigna unguiculata* and *P. vulgaris* (Holbrook et al. 1994; Fig. 2c). *G. max* is tetraploid, having an additional ancestral genome duplication event compared to related beans (Yuan and Song 2023). This higher dark inhibition variation may be explained in part by greater genotypic variation in production and regulation of CA1P. Chloroplast level differences in Rubisco regulation by metabolites including CA1P binding affinity, and post translational modifications likely also play a role (Parry et al. 2008; Amaral et al. 2024; Lobo et al. 2024). As the dark inhibition level was calculated by the difference of Rubisco activity in dark-and light-adapted samples, the presence of daytime inhibitors could mask the true level of dark inhibition (Keys et al. 1995; Orr et al. 2023). This appears likely in some species of the Poales, Caryophyllales, and particularly some in Saxifragales (Table 1, Supplementary Fig. S3a). Interestingly, despite Fabales containing species with almost no dark inhibition, very few of these species presented negative values, suggesting low daytime inhibitor accumulation in this order.

To facilitate more accurate and thorough understanding of dark inhibition will require additional data collection, potentially via streamlined methods such as linked 3PGA-NADPH assays, however validation specifically for dark inhibition readings is needed (Sales et al. 2020). Additionally, further unpublished data for predawn and midday Rubisco activity, if available, would be a valuable contribution to broaden our knowledge of dark inhibition. For accuracy in future data collection, the presence of daytime inhibitors should be accounted for. The use of sulphate for maximal activity readings may serve as an effective control for activity measurements in the absence of inhibitors (Sage 1993; Parry et al. 2008). Focusing on key phylogenetic clades of interest, such as with *Viceae* and its relatives, we may be closer to identifying the genetic controls for dark inhibition level variation.

For low inhibition species, trends in photosynthesis type were clearer than geographic trends. The majority of C4 species for which data is available had little to no dark inhibition with the most inhibited species showing <40% dark inhibition (Table 1). C2 species and species with combined CAM and C4 photosynthesis types also had low inhibition (Table 1). Though data is limited, this trend suggests a potential correlation of low dark inhibition with C4-type photosynthetic adaptations in plants (Sage and Seemann 1993). As the evolution of C4 photosynthesis often requires a careful balance of metabolites between cells, there may be stronger selection towards reducing and controlling the abundance of metabolites such as dark synthesized inhibitors (Schlüter and Weber 2020). While hot and dry environmental conditions are one of the recognised drivers for C4 photosynthesis, many low inhibition species are native outside of these conditions and are found across many environments (Stevens 2025). Even with the somewhat limited data currently available, shifts towards low dark inhibition do not appear likely to be explainable by only climate differences. For phylogenetic trends, the low inhibition tribe *Viceae*, with Mediterranean origins, is closely related to the higher inhibition tribe *Trifoleae*, suggesting a nearby ancestral dark inhibition loss event (Table 1; Supplementary Fig. 3b; Holbrook et al. 1992; Schaefer et al. 2012).

In contrast to low inhibition, the low occurrence of ‘very high’ dark inhibition suggests potential specific niches for these species. This group included CAM species (*Ananas comosus*, *Bromelia pinguin, Neltuma julifora, Clusia fluminensis*), an aquatic lily (*Nuphar lutea*), and several species in the genera *Phaseolus*, including common bean (*P. vulgaris*). No immediate common link between these species by morphology or lineage was obvious. Nearly all very high inhibition species are found in the Americas, more specifically in Central and South America, however *N. lutea* is native to Eurasia (Stevens 2025). An evolutionary pressure to retain higher levels of dark inhibition in tropical environments has been suggested previously in literature, and this trend across otherwise disparate lineages would potentially support this (Holbrook 1992). Given the high proportion of CAM species with very high dark inhibition -- six of eight non-C4 CAM species having ‘high’ or greater dark inhibition levels -- there may be a link between dark inhibition and CAM-type photosynthesis (Table 1). This photosynthesis type utilizes nighttime stomatal opening, suggesting heightened Rubisco carbamylation, a condition which would favour increased CA1P binding and stronger Rubisco regulation (Holtum 2023). Theoretically, the precursor 2-carboxy-d-arabinitol as well as malic acid should both build up in the vacuole in these plants diurnally (Moore et al. 1992), being readily available for CA1P synthesis at the onset of the night period.

Suggestions for the beneficial role of dark inhibition include the protection of Rubisco against proteolysis and reducing nighttime Rubisco metabolite binding. However, neither of these theories appear to be supported by the large variation of inhibition observed, independent of species. While there is evidence that CA1P can have a protective role against proteolysis of Rubisco, and that variation of Rubisco-CA1P binding capacity may account for some of the dark inhibition variation (Khan et al. 1999; Parry et al. 2008; Parry et al. 2013), the results from this analysis do not directly support this first hypothesis. Species which are phylogenetically close would not be expected to have significant differences in Rubisco, such as *Viceae* versus *Trifoleae,* nonetheless stark differences in dark inhibition are observed (Fig. 2a; Supplementary Fig. 2b). The *in vivo* regulation of Rubisco degradation and regulation across species is still not well understood, however evidence suggests in cereal grains that this regulation by degradation is neither complex nor varies greatly across leaf age, unlike CA1P levels (Moore et al. 1995; Irving and Robinson 2005; Feller et al. 2007; Buet et al. 2019). The absence of significant quantities of CA1P in many flowering plant species suggests a less essential role in plants. It is possible CA1P could be an accessory byproduct for a non-essential ancestral metabolic pathway across plants (Moore et al. 1992; Moore and Seemann 1992). Additionally, the role of CA1P is further confounded by the presence of its phosphatase, CA1Pase, which when overexpressed in wheat— a species without significant dark accumulation of CA1P—reduces Rubisco levels and carbon assimilation (Lobo et al 2019)

The evolution of dark inhibition remains unclear even after forty years of research, though some evidence suggests dark inhibition may be more ancient than plants (MacIntyre et al. 1997; Holbrook et al. 1992). Given the presence of high inhibition in basal flowering plant branches and the presence of dark inhibition in certain algae, dark inhibition likely did not arise in plants but rather diversified within different lineages.

The analyses of Rubisco dark inhibition data available to date for flowering plants highlights both trends and conspicuous gaps in knowledge. Future analyses should consider factors that affect the chloroplast microenvironment where Rubisco resides, including geographical distribution, climate, canopy and light adaptations, plant and leaf morphology, Rubisco characteristics, and photosynthetic pathways. Identification of the *in vivo* impact of differences in dark inhibition, the core genes tied to dark inhibition regulation, and confirmation of CA1P as the main driver of dark inhibition will establish a stronger understanding of the physiological role of this enigmatic trait, with the potential to then impact crop productivity.

## Supplementary data

**Supplementary Fig. S1.** Rubisco dark inhibition data by method and by publication.

**Supplementary Fig. S2.** Determination of group levels for Rubisco dark inhibition.

**Supplementary Fig. S3.** Rubisco dark inhibition for all plants with available data.

## Acknowledgements

The authors thank colleagues in the Photosynthesis team at Lancaster University for useful discussions and data interpretation, especially Dr. Ana Lobo, Dr. Joana Amaral, and Dr. Marjorie Lundgren. We also thank Dr. Mitchell Altschuler from the University of Illinois at Urbana-Champaign for useful discussions.

## Author contributions

CNR, ECS, and DJO: conceptualisation; CNR: data curation; CNR: formal analysis; ECS, and DJO: funding acquisition; CNR: investigation; CNR, ECS, and DJO: methodology; ECS, DJO: resources; ECS, and DJO: supervision; CNR: visualisation; CNR: writing - original draft; CNR, ECS, and DJO: writing - review & editing

## Data availability

The data collated and generated for this paper is included in the excel file found in (https://github.com/cwnehls/NehlsRamos_etal_JxB_BriefComms_Code_and_Dataset). Within the GitHub file the code used for analyses and figure generation are included. The angiosperm phylogeny tree is publicly available at (https://itol.embl.de/tree/148882475326681755591567).

## Abbreviations

CA1P: 2-carboxy-d-arabinitol 1-phosphate
CAM: crassulacean acid metabolism
3-PGA: 3-phosphoglyceric acid
RuBP: ribulose 1,5-bisphosphate

